# Size dependent diffusion and dispersion of particles in mucin

**DOI:** 10.1101/2023.06.23.546333

**Authors:** Parveen Kumar, Joshua Tamayo, Ruei-Feng Shiu, Wei-Chun Chin, Arvind Gopinath

## Abstract

Mucus, composed significantly of glycosylated mucins, is a soft rheologically complex viscoelastic material lining respiratory, reproductive, and gastrointestinal tracts in mammals. Mucus may present as a gel or as a fluid, and serves as a barrier to the transport of microbes including, harmful particles, and inhaled atmospheric pollutants. Studies on mucin gels have provided insight into swelling kinetics, and the diffusion and permeability of molecular constituents such as water. The transport and dispersion of micron and sub-micron tracer particles in mucin gels and solutions differs from the motion of molecular water since the much larger tracers may interact with microstructure and larger features of the mucin network. Here, using brightfield and florescence microscopy, high speed particle-tracking, and passive microrheology, we study the thermally driven stochastic movement of 0.5 − 5.0 *µ*m tracer particles in 10% mucin solutions at neutral pH, and in 10% mucin mixed with unmodified limestone rock dust, modified limestone, and crystalline silica. Particle trajectories, mean square displacements, and the displacement probability distributions, are used to assess tracer diffusion and transport. Complex moduli are concomitantly extracted using microrheology techniques. We find that under the conditions analyzed in our experiments the mucin solution present as a highly viscous, weakly viscoelastic fluid rather than as a viscoelastic gel. For small to moderately sized tracers with diameter than 2 *µ*m, effective diffusion coefficients follow predictions of classical Stokes-Einstein relationship. Diffusivity in rock-dust laden mucin is surprisingly larger than in bare mucin. Probability distributions of squared particle displacements indicate that heterogeneity, transient trapping, and electrostatic interactions impact tracer transport, especially for larger tracers. Our results prompt further exploration of physiochemical and rheological mechanisms mediating particle transport in soft, viscoelastic biopolymer networks and materials.

## I. INTRODUCTION

Mucus is a soft viscoelastic material that lines the surface of respiratory, reproductive, and gasterointestinal tracts in mammals. In the human body, mucus serves a multitude of crucial functions that include acting as a lubricating agent, and serving as a highly selective and permeable barrier to the infiltration and transport of pathogens and particulates to epithelial layers [1–5]. Composed primarily of water (∼90-95%), a primary constituent of mucus is mucin, polymeric glycoprotein macromolecules that contribute to its viscoelasticity and stickiness. Mucus may exist as a highly viscous fluid, a viscoelastic fluid, or as a viscoelastic gel depending on its composition, temperature, and pH. Bulk rheology measurements in the linear viscoelastic regime of gastric mucin suggests that a sol-gel transition from a viscoelastic solution at neutral pH to a soft viscoelastic gel occurs in acidic conditions beyond a critical pH of around 4 [4]. The gel-like form arises due to network formation via disulfide cross-linking; mesh sizes typically range from 200-400 nm [6, 7]. The non-linear and complex rheology of mucus also includes shear thinning properties especially in respiratory diseases; in this state, the decreased fluidity of mucin leads to congestion in lungs and may even cause bacterial overgrowth [8].

Aside from altered properties during infection, the mechanical properties of mucus, and its response to flow and deformation, is strongly impacted by embedded colloids and inhaled environmental particulates and aerosolized chemicals. For instance, nicotine, the active ingredient in cigarettes and e-cigarettes, has been found to increase mucus viscosity significantly, even at low-concentrations [9]. In the mining industry, limestone dust is often utilized as a safety measure against coal dust explosions. Exposure to such dust particles results in serious risks to health as the inhaled particulates can affect mucus rheology and swelling, amongst other biophysical properties [10–12]. Indeed, respiratory diseases as a result of working in mines, such as chronic obstructive pulmonary disease (COPD), is linked with changes to mucus rheology [10, 11, 13, 14]. Motivated by this, many recent studies on unmodified limestone (UCRD), and surface-modified limestone (MTRD), have explored changes in viscosity, mucus permeability, and diffusion of water associated with swelling (for instance see [15], and references therein). Mucus is commonly characterized using macrorheology measurements that allow the determination of bulk, macroscale properties such as shear viscosity, and effective dynamic loss (viscous) and storage (elastic) shear moduli [16, 17]. For effective function, these properties are needed to lie within physiologically determined ranges [3].

The barrier properties of mucus and mucin in the body has motivated many recent studies on reconstituted mucin and synthetic mucin based biopolymers and biogels. Commercially available mucins do not form hydrogels in physiological conditions, and the process to purify them are laborious and relatively low-yielding [18]. The gelation of mucin polymers are affected by the composition of the mucin chains themselves such as type, purity, and length. Furthermore, the physiological environment such as pH, ionic strength, and the presence of calcium ions can affect the ability of mucin to maintain its viscoelastic properties [19]. Thus significant challenges remain in realizing engineered polymeric materials that mimic the function and properties of mucus. More recent interest in engineering mucin-based materials arise from applications in biomedical engineering and drug delivery. For instance, mucus acts as a barrier to drug absorption efficiency by hindering nanomaterial transport; however, the use of carboxymethyl cellulose (CMC) additives has been shown to improve drug absorption efficacy [20, 21]. Carboxymethyl cellulose (CMC) is a cellulose-derived polymer comprised of long polymer chains. Spanning a range of molecular weights from a few kilo-Daltons to mega-Daltons, CMC is ubiquitously used in cosmetic, food, and pharmaceutical industries [22, 23]. Solutions of CMC comprise of entangled polymers and porous biopolymer networks that - like mucin solutions and gels - enabling selective passage of macromolecules. The ability for CMC to have tunable viscoelastic behavior [23, 24] makes the widely available product a prime candidate for exploring its use in mucin-based composites.

While bulk swelling measurements have been conducted on mucin gels, and mucin-dust systems, the thermally driven random motion and transport of particulates with diameters from sub-micron and micron values (sizes typically encountered as atmospheric aerosols) is largely unexplored. Particles sample mucin microstructure at a range of length scales in a size dependent fashion. Thus small particles will sense and respond to the mucin environment differently than large particles. As a result, overall particle diffusion, dispersion, and penetration will be impacted by strongly by microscale heterogeneities and therefore cannot be explored solely using bulk rheology measurements.

Techniques that enable direct visualization of tracer particles embedded in mucin and track their motion in real time are well suited to explore these questions. Microcopy based particle-tracking methods, and the analysis of tracking data with microrheology theory are well-established, and have been used with great success in exploring the rheological environment of soft materials including complex fluids and gels, [25–29]. Typically image-based tracking passive microrheology observes these particles undergo Brownian motion with either brightfield or fluorescence microscopy where subsequent images are taken at high frame rates. The generalized Stokes-Einstein relationship in the complex domain can be then used to relate the mean-squared displacement (MSD) of the tracer particles to the dynamic shear moduli of the ambient medium [5, 27, 30–32]. We have also recently used particle and cell tracking to understand local fluid flows and the rheological environment of active and biological materials including bacterial suspensions [31, 33, 34], bacteria swarms [35], algal suspensions [36]. Particle tracking combined with traction force microscopy has also been used to extract viscoelastic and elastic material properties that are needed in computational and theoretical studies of biological matter [37].

Here, we use brightfield and florescence microscopy, high speed particle-tracking, and passive microrheology, to study the diffusion, transport and trapping of spherical tracer particles in reconstituted mucin and mucin-laden with various types of commonly used rock dust. The organization of the paper is as follows. In section §2, we describe the methodology and summarize the theoretical underpinnings of the analysis. We then study the diffusion of 0.5 − 5.0 *µ*m tracer particles in Newtonian DI water, and in a model viscoelastic CMC solutions. Having validated the methods and analysis, in Section §3, we present our analysis of tracer transport in reconstituted bare and dust-laden 10%mucin. We conclude by suggesting physical mechanisms to explain observed particle dispersion data, and discuss how tracer size combined with mucin properties impact tracer transport.

## II. MATERIALS AND METHODS

### A. Preparation of control mucin and mucin loaded with anti-caking agent (Rock dust)

The rock dust types used our study were: (1) unmodified limestone rock dust (MineBriteTM G; UCRD with mean particle size ≤74 *µ*m), (2) modified limestone which is a moisture-tolerant rock dust, MTRD and has mean particle size 19.5 *µ*m), and (3) crystalline silica (Min-U-Sil®10, SiO_2_ with mean particle size 3.4 *µ*m). Mucin samples (Sigma-Aldrich, Type III Mucin from porcine stomach, M1778) were prepared by mixing mucin granules with PBS to a final 10 wt% concentration (at pH = 7.3). This was set as our control system. Rock dust solutions were prepared by dissolving particles in Hank’s buffer for 24 hours to a concentration of 1 mg/mL. The final formulations were: (a) (Mucin-CD) mucin with UCRD in solution, constituted by mixing 1 mg/ml of UCRD and 10 wt% mucin in DI water; (b) (Mucin-CD) mucin with MTRD in solution, constituted by mixing 1 mg/ml of MCRD and 10 wt% mucin in DI water; (Mucin-S) mucin with SiO_2_ in solution, constituted by mixing 1 mg/ml of SiO_2_ with 10 wt% mucin in DI water. We did not vary pH, or add crosslinker to our reconstituted mucin formulations. Based on prior work, we anticipate that our mucin samples will behave as viscoelastic solutions as opposed to viscoelastic gels. Visual inspection confirmed with these expectations.

### B. CMC Sample Preparation

We used carboxymethylcellulose (CMC) from Sigma-Aldrich (MW = 250 kDa and DS = 0.7), although solutions with MW = 90 kDa were also tested. Homogeneous solutions with 0.5%, 1%, and 2% (in weight %) concentrations of CMC were formulated. CMC samples were prepared at 40°C and spun on a magnetic stir plate at 100 RPM for 48 hours. Samples were then allowed to rest for 1 hour prior to measurements to allow the network to relax.

### C. Slide Preparation for particle tracking and microrheology

Imaging channels were designed with McMaster-Carr polyester plastic mounting tape (Product No. 75955A673) by folding the tape onto itself, and subsequent smoothing. This resulted in a well geometry that was ≈200 *µ*m in depth. The imaging well was punched out from the folded tape with a 0.5 inch in diameter hole-punch and then affixed to a glass slide of dimensions, 25 × 75 × 1 mm (Fisher Scientific). For imaging experiments, 16*µ*L of sample solution was first pipetted into the well, then a cover glass slip (18 × 18 mm, 0.13 mm thickness, VWR) was placed over the sample and fixed by double-sided mounting tape. The tracer particles were diluted to 1:100 concentration in DI water and injected into the samples. The injected volume was small so that local water content remained approximately the same.

### D. Optical Setup for microrheology

For particle tracking and microrheology measurements, we used spherical florescent particles of diameter *a* = 0.5−5 *µ*m (Spherotech, Nile Red, Excitation wavelength, *λ* = 510 nm), and also non-florescent Spherotech polystyrene particles of diameter *a* = 0.5 − 5 *µ*m as tracers. A Zeiss 200m Axiovert microscope in brightfield mode (for larger tracers), and sometimes in fluorescence mode (for smaller sub-micron particles) was used to deliver high contrast images. Post loading, samples were allowed to equilibrate and data was recorded after 10 minutes. The motion of the tracer particles are subsequently captured for 1 to 10 minutes and are saved as AVI videos.

All images presented were taken with a Zeiss EC Plan-Neofluar 40x/NA 0.75 M27 (Working distance = 0.71 mm, Depth of Field = 1.09 *µ*m) objective. Video was recorded using a Mako G-158B monochrome camera. Particles in mucin samples were imaged at 30 fps with 30 ms exposure time. Particles in CMC were imaged at 90 fps with 11 ms exposure. Video recordings were done with the imaging plane focussed on the center plane of the rectangular channel (≈ 100*µ*m away) to minimize hydrodynamic, surface, and capillary effects from the channel edge walls. The experiments were conducted at room temperature (21°C) measured with a Neulog temperature sensor (NUL-203).

### E. Image filtering and pre-tracking processing

Image filtering and particle tracking are based off of Crocker and Grier’s particle tracking algorithm adapted to MATLAB [38]. To accurately reconstruct trajectories from image stacks, the particles should be clearly visible in the image and well contrasted against the background so that the tracking algorithm can easily detect those particles and link detected particles between frames while discriminating between particles. Movies and associated TIFF stacks were batch processed to main consistent contrast within each video.

Figure 1 illustrates the raw images of particles from experiments conducted using brightfield mode (Figure 1(I-III), top three tiles from left to right), and for experiments using florescent tracers (Figure 1(I-III), bottom three tiles from left to right). Images of individual tracer particles display concentric rings around them with decaying intensity (see the closeup in Figure 1(III), bottom). The central ring of this diffraction pattern, ie., the Airy ring, has the highest intensity and was used to fit and calculate particle locations. The intensity profile of the Airy ring is approximated by the form [38, 39],

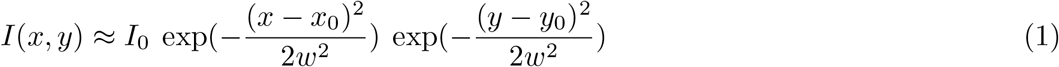

where *I*_0_ is the peak intensity (amplitude) at the center, *x*_0_ and *y*_0_ are the coordinates of the center, and *w* is the root mean square width. The resolution of images is limited by the Rayleigh criterion. Two blobs (intensity fields associated with two particles) separated by more than diffraction limited distance could be resolved as two separate entities. If the center-to-center distance of the Airy rings is less than the diffraction limit, they cannot be resolved as two separate entities. Since the centroid positions are determined by fitting the intensity profile function, a resolution of a few nanometers is possible.

**FIG. 1.**
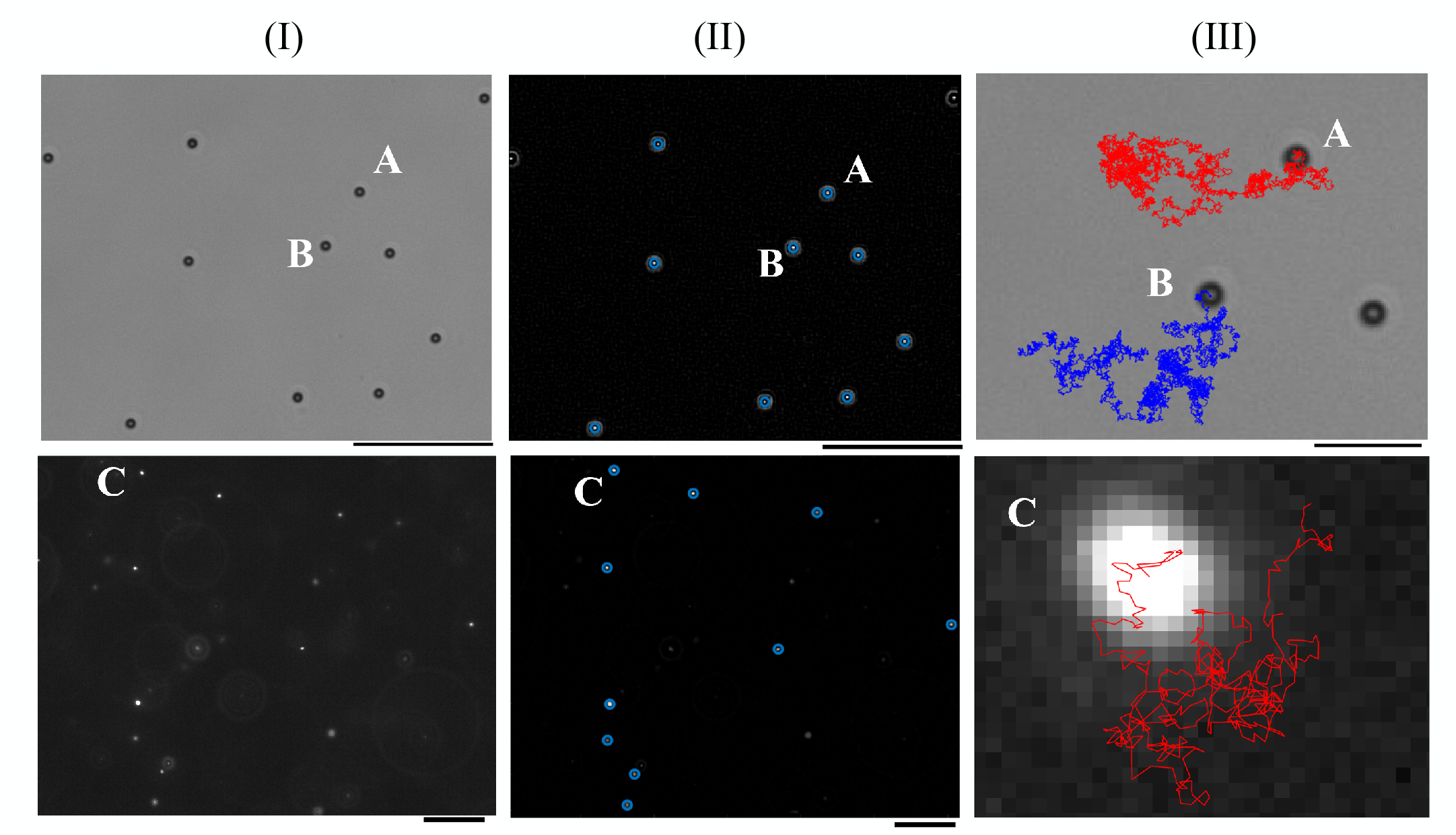
Spatiotemporally resolved particle tracking using brightfield microscopy for larger tracers, and florescence microscopy for sub-micron tracers. We show the three main steps in stitching and composing particle trajectory videos and image stacks from raw data. Trajectories are computed in the *x* − *y* plane corresponding to the imaging focal plane. The top row illustrates the method for brightfield images, and the bottom row for images obtained using florescence. For each case, columns depict (I) tracer particle detection, (II) image filtering, and contrasting, and (III) trajectory construction. **(I, top)** The original image of 2.29 *µm* Spherotech polystyrene tracer particles in DI water (Zeiss 200M Axiovert microscope, 40x/NA 0.75 objective, 30 fps, 30 ms exposure time. The total observation time = 600 s and temperature = 21°C). **(II, top)** Filtered image of (I, top) with automatically identified particles, following the Crocker-Grier algorithm. Bandpass filters ensure that particles are shown as bright spots against a dark background. **(III, top)** Two representative tracks from the particle tracking routine. **(I, bottom)** Original florescence microscopy image of 0.53 *µm* Spherotech fluorescent tracer particles in 90 kDa (CMC) solution (Zeiss 200M Axiovert microscope, 40x/NA 0.75 objective, 90 fps, 11 ms exposure time. Total observation time = 60 s). **(II, bottom)** Filtered image of (I, bottom). **(III, bottom)** Magnified image showing only trajectories of the particle identified as C.

There exist many algorithms in literature to determine the centroid positions of the tracers; here we use the well-established algorithm due to Crocker and Grier [38]. In order to identify and determine the center of each tracer particle, recorded images extracted from the videos were prepared to remove the background and other artifacts. The contrast was adjusted so that the particles are as bright as possible against the background and the image was converted to greyscale. Spatial bandpass filtering was used to remove spurious features, and any bright spots present which did not fall within the particle size limits. The centroids of the remaining bright features were then determined. Spurious extreme particle sizes resulting from imaging two or more particles adjacent to one another were rejected in the tracer count. Even after employing the bandpass filter, some artifacts (non-tracers) remained as faint spots that were detected as real tracers by the algorithm. To remove these artifacts, images were filtered so that only particles above a brightness threshold remain were identified as real tracers. An example of this is illustrated in Figure 1. Specifically, in Figure 1(II)top and bottom, we see real tracers particles identified (in one frame) and marked with a blue circle, and these are recorded in a database with a unique identifier.

Trajectory construction is done by finding a tracer particle in a given image and connecting to the most likely corresponding particle in the succeeding image. The algorithm adopted takes into consideration the dynamics of the non-interacting Brownian motion of particles. For interacting particles, the video is recorded at high frame rates and a minimum squared displacement threshold is defined such as particles below that threshold are removed from the analysis (when particles stuck temporally as doublets). Finally, a memory function is defined to account for particles moving in-and-out of the focal plane. The memory function defines a maximum amount of frames an already detected particle may lose detection and re-gain detection. For example, a detected particle may leave the focal plane at time *t*_0_ for 2 frames due to thermal noise and will not be detected for those 2 frames, but be detected in the subsequent frame (*t*_3_); the memory function will declare that the particle at *t*_0_ and at *t*_3_ must be the same particle and will link the two detection spots together to generate a single track. We manually inspected many of these tracks to confirm the analysis. Stitching together the positions of such identified particles provides the coordinates of the tracers in time and allows us to reconstruct the raw trajectories (the blue and the red curves) as shown in Figure 1(III). The memory feature of the MATLAB function we use also defines the number of frames a particle can be lost before considering subsequent emergence and motion as as a new track.

Finally the raw trajectories were corrected for drift using a statistical model included in the particle tracking algorithm that employs velocity drift corrections (see Figure 2). Drift arises from various sources including small amplitude stage movement, sample leakage, building vibration, and thermal noise due to heating. We did used vibration-free stages to minimize the vibration induced drift so that these biases were reduced.

**FIG. 2.**
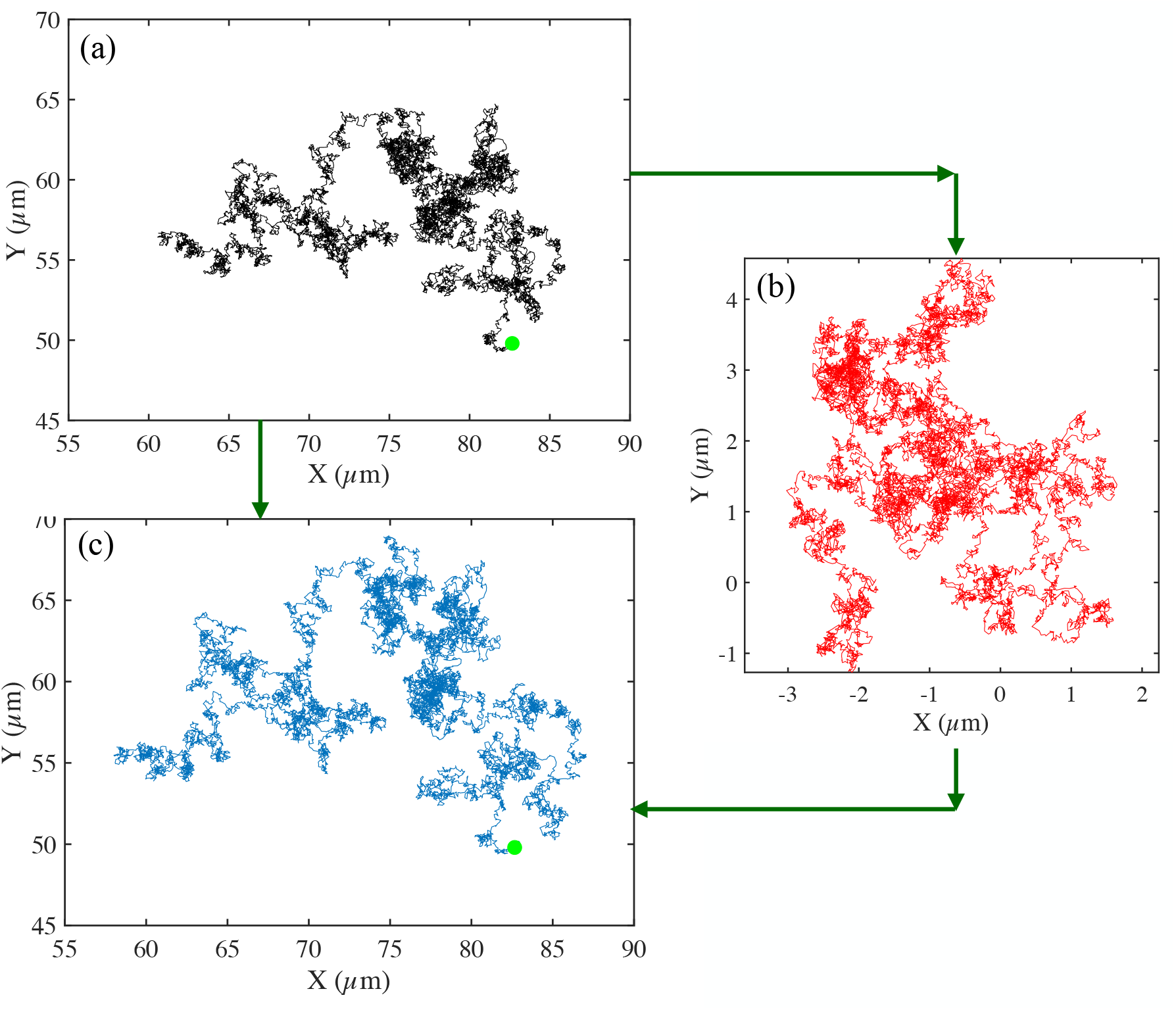
Example raw trajectory corrected for drift to obtain the final reconstructed trajectory. Trajectories are computed in the *x* − *y* plane corresponding to the imaging plane. We show trajectory construction, and drift correction for the motion of a 2.29 *µm* tracer particle in DI water (from Figure 1, top) is shown. In **(a)**, we show the uncorrected particle trajectory, with the start of track indicated by the green circle. In **(b)**, the drift vectors are plotted for the full observation time (600 s). Here, drift acts as undesired translation of the particle, affecting its natural trajectory through a medium. In **(c)**, we plot the corrected track (trajectory) with drift subtracted out from the original trajectory. The start of track is indicated with the green circle. Images were taken on Zeiss 200M Axiovert microscope with 40x/NA 0.75 objective at 30 fps, 30 ms exposure time, and temperature = 21^*o*^*C*.

### F. Statistical Analysis

The ensemble averaged mean square displacement of tracers - that is, an average over a population of isolated tracer particles - as a function of delay time τ (ie., ⟨MSD(τ)⟩ is defined by

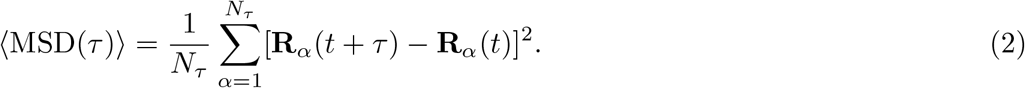

Here, **R**_*α*_(*t*) is the reference position (in *x y* cordinates) of tracer with index *α* at some time *t* along its trajectory, **R**_*α*_(*t* + τ) is position of the same tracer at time *t* + τ, and *N*_τ_ is the total number of particles (individual trajectories). As tracer particles can leave the imaging plane, not all particles have the same value of the maximum delay time.

It is important to clarify that there are two types of averaging employed in Equation (2). First is at the level of each tracer trajectory generated over an observation time *t*_*F*_. Fixing a time delay τ, we calculate the square displacement for all possible pairs of time instances separated by τ and calculate an average MSD from these values. This MSD is then generated for all possible delay times along the full single trajectory to generate MSD(τ). A second level of averaging comes from taking an ensemble average of multiple tracer particles to then generate ⟨ MSD(τ) ⟩. Note that small delay times will have a larger number of trajectories while fewer trajectories are associated with large delay times. Hence statistical error is expected to be larger for larger delay times.

Ensemble averaging hides possible spatiotemporal inhomogeneities that are associated with the ambient environment, in this case the mucin and CMC formulations. To gain more insight about how each tracer explores its local environment, we analyze discrete probability distributions (histograms) of tracer displacements. These histograms are obtained on a particle (or trajectory basis) and are averaged over the trajectory of a tracer *but are not averaged over particles/tracers*. Thus, they provide a measure of variations in the local environment sensed by each particle.

For each trajectory, we divide the overall displacement time history (from time *t* = 0 to the final time for which the trajectory exists) into intervals of τ. We then compute the squared displacement between reference times *t*_*R*_ and time *t*_*R*_ + τ repeating this exercise for all possible values of *t*_*R*_. Thus by averaging over *t*_*R*_, we obtain the trajectory averaged mean square displacement *for a single tracer* as a function of the delay time τ. We then bin the results using bin-widths of 0.05 *µ*m^2^ and generate a histogram using

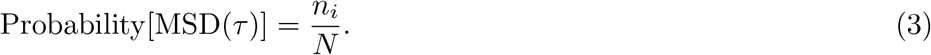

Here, *N* is the total number of samples and *n*_*i*_ is the number of estimated samples (the number of MSD values) within a bin. Note that via this calculation, each trajectory (tracer particle) is assigned a mean value of a squared displacement for a delay time τ. The histogram merely provides the probability distribution from the many trajectories. Values and histograms are generated for 3 time increments τ = 0.1, 1, and 10 s to assess the motility of particles within the networks at short and long time scales. All errors bars correspond to 1 standard deviation.

### G. Velocity correlation, diffusion, and dispersion of tracers

Following an ensemble of equally sized tracers, we first obtained the probability distribution (histogram) of displacements in a fixed time interval (delay time). For freely moving, and non-interacting Brownian particles, the ensemble average of displacement of is expected to be zero, and Gaussian with width determined by the tracer diffusivity and the delay time). Figure 3a shows one such histogram (a discrete version of the probability distribution function) for 2.29 *µ*m particles (Figure 3, a) moving in DI water at 21°C. Here the *x* and *y* coordinates are measured in a fixed lab-frame and span the focal plane. As expected, the *x* and *y* displacement histograms of particles are similar since there is no directional bias to the motion that breaks *x* − *y* symmetry. Both distributions are Gaussian with zero mean and identical width - here dependent on the diffusivity and the time increment over which the displacements are made.

**FIG. 3.**
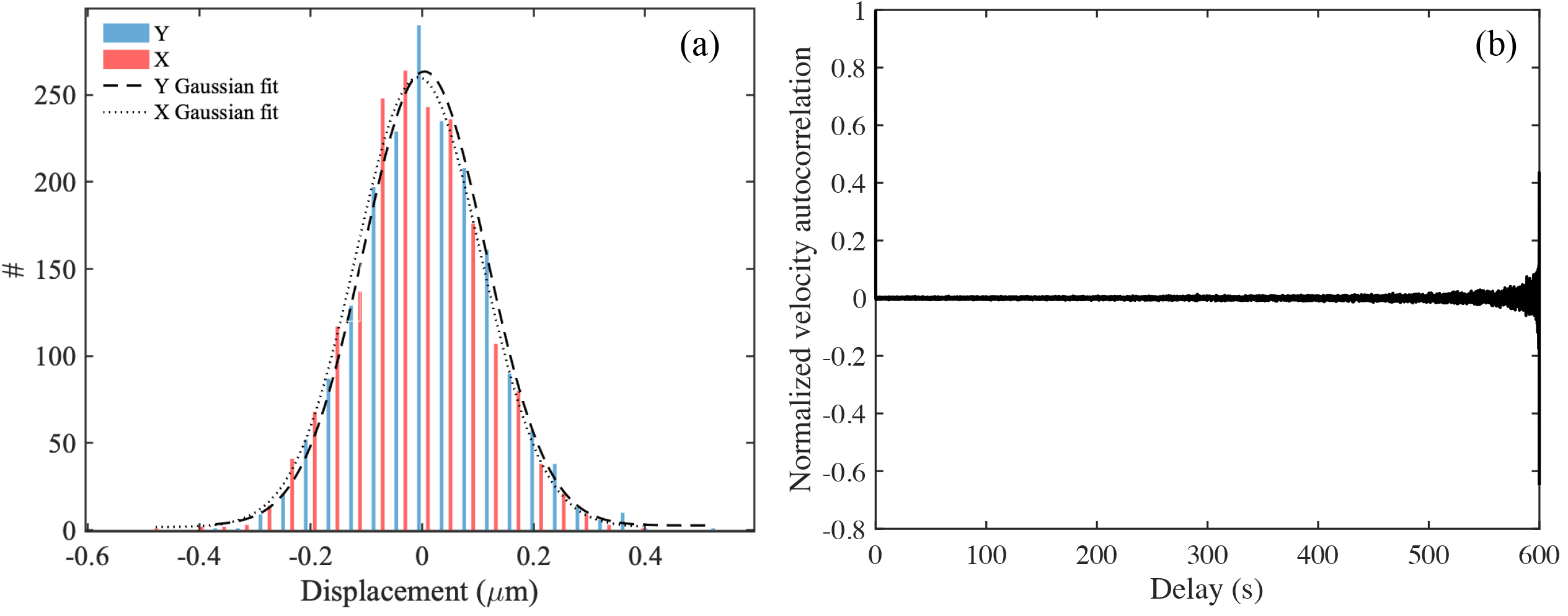
Analysis of tracer particle trajectories of the 2.29 *µ*m tracer particles in DI water, a Newtonian fluid. **(a)** Histogram of (particle averaged, over single particle trajectories) translational displacements in *x* and *y* Cartesian coordinates as tracked from Figure 1, displaying a Gaussian distribution with zero mean and finite variance (equal for both *x* and *y* components) for the probability distribution of tracer displacements. Since particles undergo free Brownian motion, the variance is related to the diffusivity and the delay time over which the displacements are evaluated [40]). **(b)** The normalized velocity autocorrelation function as a function of delay time (in seconds) for trajectories recorded. The velocity autocorrelation is nearly zero, confirming the Brownian motion of the tracers in DI water. Large fluctuations for large delay times are due to a decreasing number of sample trajectories.

The velocity of diffusing (Brownian) tracer becomes uncorrelated with prior values as the moving particle moves along its trajectory in time and loses memory of its prior spatiotemporal position. This may be quantified using the velocity autocorrelation function applied to a single tracer particle,

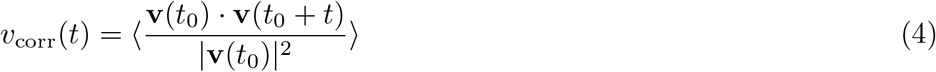

where *t*_0_ is a suitable chosen initial reference time. The velocity correlation for 2.29*µm* polystyrene particles in water is shown Figure 3b. A value of zero shows that the particle motion is uncorrelated and confirms a random walk motion where. The curve is normalized so that we get value of 1 for *dt* = 0.

In two dimensions as is appropriate in our experiments in DI water, the trajectory-averaged MSD of Brownian tracers valid in the long time limit, and the ensemble averaged MSD are statistically identical with the same mean. This mean is related to the particle diffusivity *D* and time *t* by the relationship (see for instance, [40]):

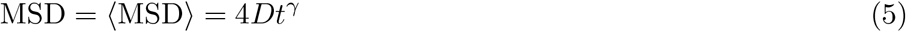

with *γ* = 1. Sub-diffusion is characterized by *γ* < 1 while superdiffusion corresponds to motion with *γ* > 1. The diffusivity *D* (in Equation 5) for spherical tracers of diameter *a* moving in a Newtonian fluid at viscosity *µ* and temperature *T* is given by the Stokes-Einstein relationship

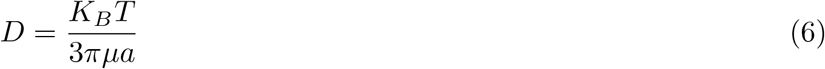

where *k*_*B*_ is the Boltzmann constant. Figure 4a shows individual tracer mean square displacement values calculated for each independent trajectory/track for 2.29 *µ*m tracers in DI water. Ensemble averaging the values we obtain a linear relationship between ⟨ MSD ⟩ and *t* confirming the freely diffusing motion of the particles. Note that the slope of the curve (or equivalently the value of the ensemble averaged MSD evaluated at some time *t* = τ can be used to estimate *D* for this particular tracer size. Alternately, knowing the temperature *T* and tracer diameter *a*, one can use Equation 6 to calculate the effective viscosity of the ambient medium.

**FIG. 4.**
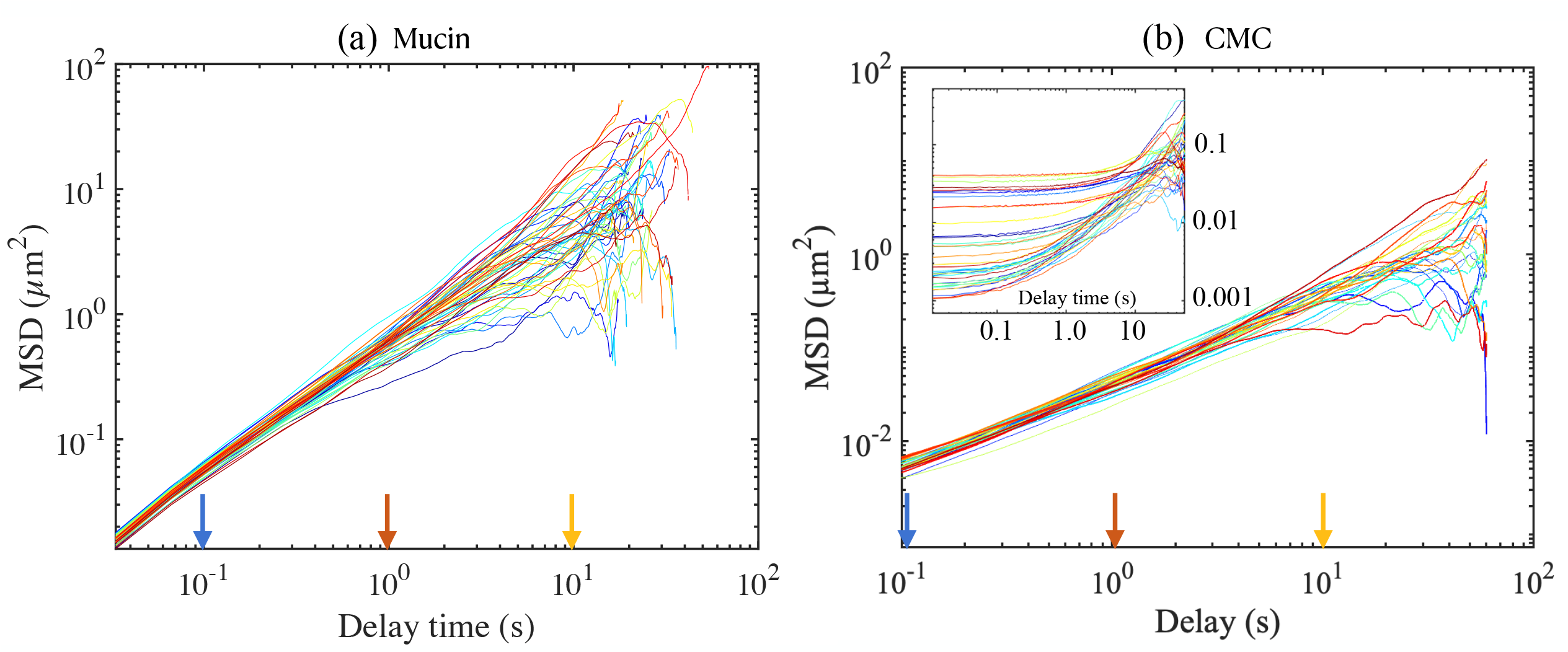
Single particle Mean Square Displacement (MSD) of tracers as a function of delay time for (a) unloaded 10 wt% mucin solution, and (b) a solution of CMC (concentration 1 wt%, MW 250 kDa). Tracer sizes are (a) 1 *µ*m, and (b) 0.87 *µ*m, and the number of tracks are (a) 157 and (b) 37, and for each the ensemble averaged value ⟨ MSD ⟩ (τ) is calculated by averaging over the values at each τ. The power law exponent *γ* in the relationship MSD ∝ τ ^*γ*^ is 2 for a ballistically moving tracer, *γ* = 1 for freely diffusing tracer and *γ <* 1 for a sub-diffusing tracer. Also indicated are the three values of the delay time τ = 0.1s, 1s and 10. Temperature = 21°C. The inset in (b) shows typical MSD’s for a more concentrated 2% CMC solution obtained for a larger 2.11 *µ*m tracer.

### H. Diffusion of tracers in viscoelastic CMC

The mean square displacements statistics of located tracks can be transformed to calculate the frequency dependent linear viscoelastic moduli of the ambient fluid and assess its elastic behavior and quantify its rheology.

Figure 4b shows sample MSD curves evaluated from trajectories of individual tracers in CMC solution, a canonical viscoelastic fluid at concentrations. The CMC formulations correspond to solutions that are 0.5, 1, and 2% (weight/volume) and thus are expected to be increasingly viscoelastic. Tracers used were 0.87*µ*m, and 2.11 *µ*m diameter particles. A dramatic difference is seen in the main figure tile, compared to the inset. The difference may be attributed to the viscoelastic behavior of the CMC at the higher concentration *and* to the sampling of larger network pore sizes and interaction with the entangled polymer network by the larger tracer particle.

Ensemble averaged curves MSD(τ) can be converted to appropriate MSD in the Laplace (Fourier) frequency domains (note that 1*/τ* where τ is the lag time, can be interpreted as frequency). Application of the generalized Stokes-Einstein relationship (GSER) [29, 41, 42] then provides the complex moduli in the frequency domain. The GSER is built on the principle of average motion of tracer particles in a continuum complex fluid, and is therefore valid provided macrostructural features are smaller than the tracer sizes. The frequency dependent version of the Stokes relationship effectively provides a measure of the viscoelastic drag on the tracer particles. Assuming local homogeneity and isotropy, this may be analyzed to obtain the complex shear modulus of the ambient medium.

The viscoelastic modulus 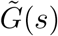 calculated from the unilateral Laplace transform of ensemble averaged 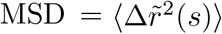 is given by

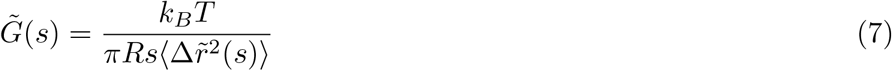

where *k*_*B*_ is the Boltzmann’s constant, *T* is temperature, *R* is tracer particle radius, and *s* = *iω* is Laplace frequency [27, 28, 42]. Following previously established theory, ⟨Δ*r*^2^(*t*)⟩ is expanded algebraically in a power law, and leading terms are retained to calculate the viscoelastic moduli. We use

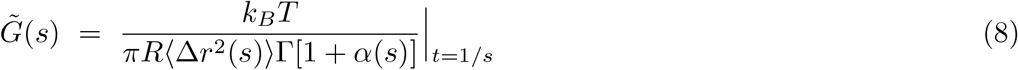

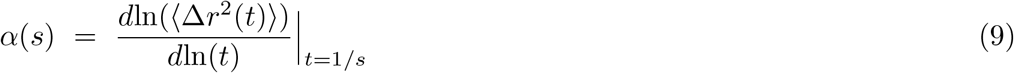

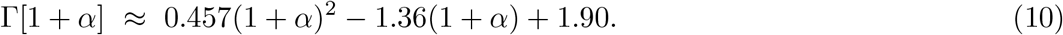

The storage (elastic) modulus *G*′, and the loss (viscous) modulus *G*′′ may are obtained from Equation 8 by extracting the real and imaginary components respectively. If *α* ≈1, we expect the tracer particle to be moving diffusively in isotropic Newtonian fluids. For tracer particles surrounded in a purely elastic medium that severely inhibits their thermal motion, *α* ≈ 0 and the storage modulus *G*′ is the leading term in the complex modulus.

## III. RESULTS AND DISCUSSION

### A. Reconstituted mucin solutions behaves as very weakly viscoelastic fluids

Previous studies using commercially available mucin have found that reconstituting and hydrated the industrially produced powder typically yields solutions rather than gels. The solutions are highly viscous with weak elasticity. The lack of gelation is likely due to manufacturing methods [17, 43]. Adding chemicals that enables crosslinking allows gelation of commercial mucin, but is laborious and requires additional extensive characterization [18]. Since we do not add crosslinkers, we expect our10 wt% mucin solutions to behave as mainly viscous fluids. Indeed, averaging over the MSD obtained from individual trajectories for tracers moving in the bare 10% mucin solution, we find that the ensemble averaged MSD scales with time as *t* (exponent *γ* ≈ 1) for lag times approximately less than τ ≤ 8 s, and sub-diffusive behavior with *γ* < 1 for *τ* > 8 s. The cut off time is subject to statistical error since the number of observed trajectories decreases with larger lag times. The cut-off value also depends on the tracer diameter. Tracers with diameters 2.11 *µ*m and smaller followed the diffusive behavior for longer lag times, while the 5 *µ*m tracers deviated at shorter lag times.

The MSD’s obtained for the 10 wt% mucin was very different from that seen for the 2% CMC solution (Figure 4(b) inset); difference here highlight how viscoelastic effects modify and impact the forms for the MSD’s at small delay times. These results were consistent with our microrheology analysis. We extracted the complex modulus *G*^*^ = *G*′(*ω*) + i*G*′′(*ω*) from ensemble averaged MSD data for both mucin and CMC solutions and found different behaviors. For the 10 wt% mucin solution, we found that *G*′′(*ω*) was the dominant component and tracked | *G*^*^ | very closely. The elastic component *G*′(*ω*) ≪ *G*′′(*ω*) for frequencies less than around 30 Hz, and while increasing as a function of frequency thereafter, was still less than the loss modulus. Figure 5(a) shows the magnitude of the complex modulus as a function of the frequency obtained using particle tracking data.

**FIG. 5.**
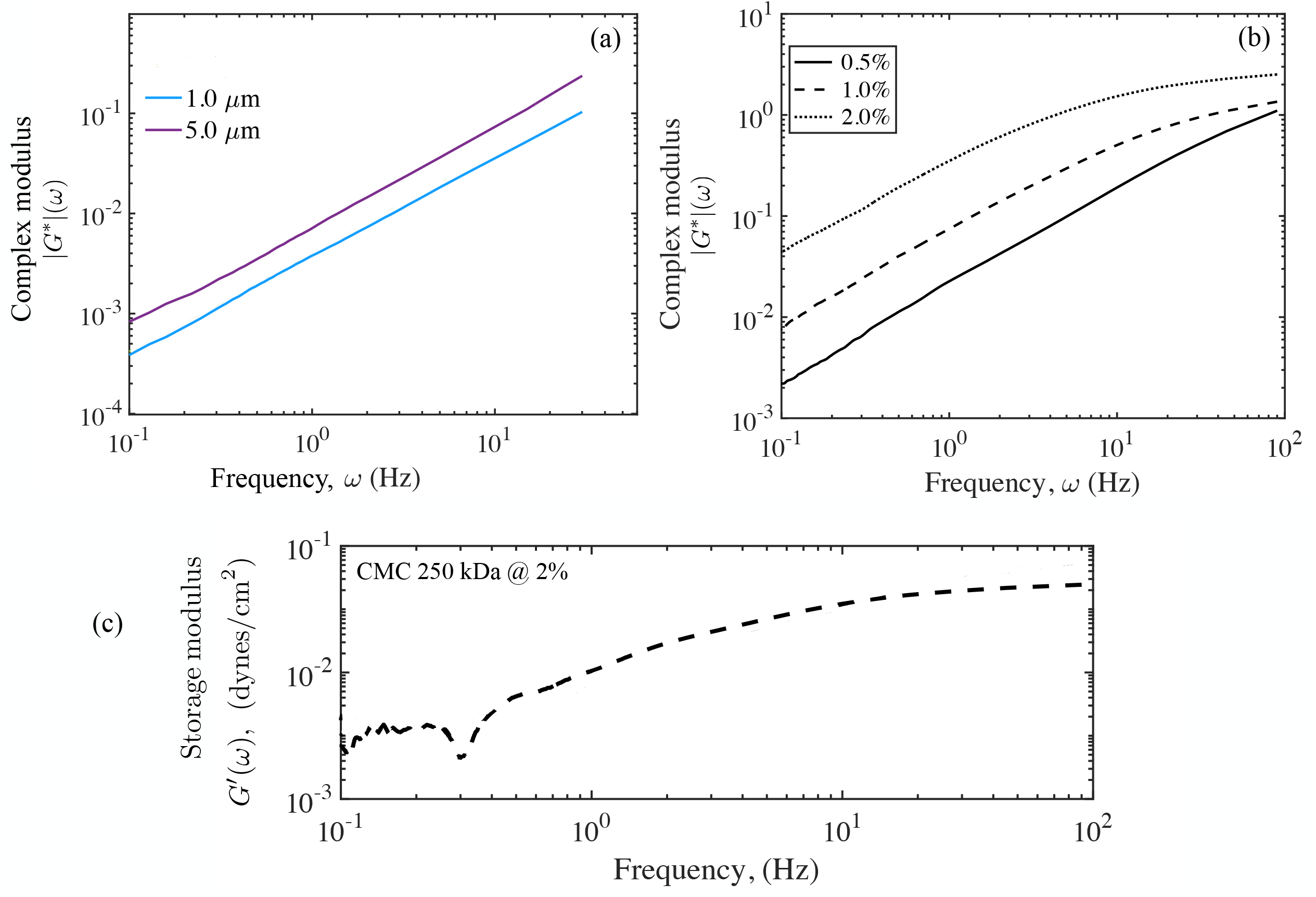
Magnitude of the complex moduli, |*G*^*^(*ω*) |, of the unloaded 10 wt% mucin (control sample), and of solutions of 250 kDa CMC at varying concentrations obtained using microrheology. **(a)** The Generalized Stokes-Einstein formulation was used to calculate the complex (*G*^*^) moduli and its magnitude of control mucin using tracer sizes of 1.0 *µm* (blue), and 5.0 *µm* (maroon). For the small to moderate frequencies *ω* shown here, a nearly linear behavior is seen with frequency for both, indicating that the mucin solution is dominantly viscous. **(b)** Microrheology results for 250 kDa CMC solution at 0.5, 1.0, and 2.0% (w/v) for 0.87 *µm* tracers are shown here. The storage modulus *G*′(*ω*) and the loss modulus *G*′′(*ω*) were also estimated separately for the CMC solutions, with significant elastic components suggesting strong viscoelastic response unlike for the 10% mucin solution. (c) The elastic modulus *G*′(*ω*) obtained from analysis of trajectories for 0.87 *µ*m tracers.

To contrast the viscously dominated response with weak elasticity of the 10 wt% mucin with a more elastic response for a viscoelastic fluid, we show |*G*^*^| extracted from the appropriate ensemble averaged MSD data for CMC solutions in Figure 5(b) and *G*′(*ω*) in 5 (c). Figure 5(b) investigates the effect of CMC concentration, and illustrates the increasing elastic component as the concentration increases for all values of the frequency. Eventually at the high frequencies, we see a tapering of the curve (seen clearly for the 2% concentration) as the elastic component *G*′(*ω*) dominates over the viscous component. Increasing CMC concentration typically creates denser entangled networks yielding higher *G*′ values. We find that the complex modulus of 250 kDa CMC (G^*^) is lower than results for 700 kDa CMC [23]. Our CMC samples are with 250 kDa CMC, which consists of significantly smaller chains and crosslinks.

For 0.5% CMC, values of | *G*^*^(*ω*)| and loss modulus *G*′′(*ω*) are comparable for the the frequency ranges investigated indicating the viscous nature of fluid. For 2% CMC, the magnitude of the complex modulus | *G*^*^| is comparable to the storage modulus *G*′ of elastic gel at high frequencies. For 2% CMC, the values of | *G*^*^| track the values of *G*′′ at lower frequency and to *G*′ at higher frequency. Figure 5 (c) shows how *G*′(*ω*) varies for tracers (*a* = 0.87 *µ*m) as a function of frequency. Comparing this to 5(b), we note the clearly increasing elastic component for large frequencies. The transition for viscously dominated to elastically dominated response occurs at a characteristic frequency that is tracer size dependent.

### B. Diffusivity of tracers decreases inversely with tracer diameter

To study the short term transport and diffusion of tracers in the various mucin formulations, we ensemble averaged the individual MSD’s of tracer particles with varying *a* and fit the linear part to an effective tracer diffusivity *D*. To check the accuracy of the fitting routine, and the averaging of the MSD’s we first analyzed particle trajectories in water and obtained the diffusivity of tracers in DI water at 21°C. This data is shown in Figure 6(a) (stars connected with dotted line). Also shown as the dashed line is the theoretically expected value from Equation 6 using the viscosity of water. For smaller tracer sizes ≤1 *µm*, the experimental diffusivity is comparable to theoretical predicted values within ∼5% tolerance. Overall, we find excellent agreement for tracers ≤ 2.11 *µ*m with larger tracers showing slight deviations. We attribute this to settling out of the field of view resulting in reduced number of trajectories available for analysis and enhanced interactions with the upper and lower walls of the channel. For instance using only short trajectories with lag times less than 1s results in a larger variance (range denoted in red).

**FIG. 6.**
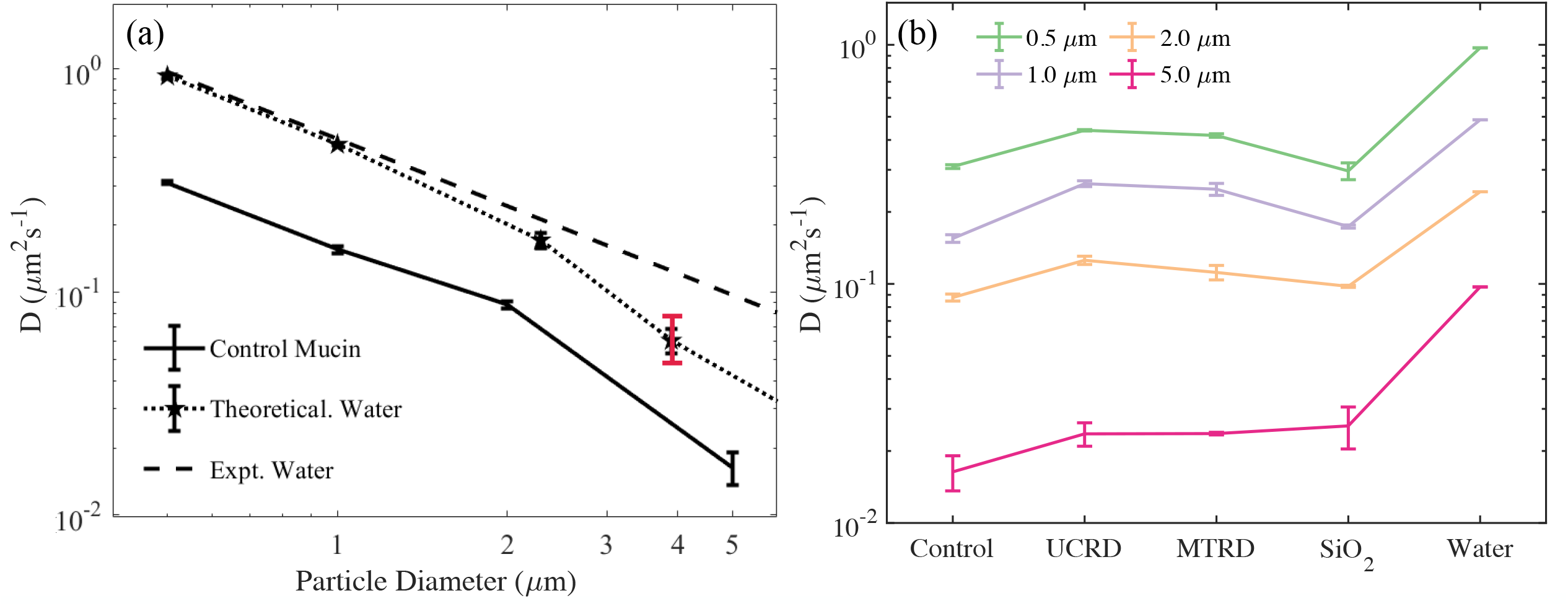
Values of the effective tracer diffusivity estimated from particle tracking and trajectory analysis. **(a)** Experimentally estimated diffusivity of tracers in DI water (data connected by the dotted line) at a temperature of 25 C. The theoretical result predicted by the Stokes Einstein relationship is shown as the dashed line. We see excellent agreement for small to micron sized tracers. For the largest tracer 4 − 5 *µ*m, we observed a decrease in frequency - this is attributed to increased sedimentation effects that resulted in fewer trajectories and a concomitant larger statistical error. Error bars (black) correspond to 1 standard deviation. The red bar indicates the variation in results when the channel size was reduced by 15 % with tracers interacting more strongly with the walls of the channel. Also plotted are the diffusivity values of tracers moving in the unloaded 10 wt% mucin solution. The control mucin solution behaves primarily as a viscous fluid for small tracer particles with an effective viscosity that is larger than DI water. Visual observations suggest that the largest tracers in these cases typically remained in the field of view but manifested a pronounced reduction in random motion **(b)** Estimated diffusivities of tracer particles 0.5-5 *µm* in diameter for the 10 wt% control mucin solution and 10 wt% mucin solution with dust and anti-caking additives. ⟨MSD⟩ for short to moderate delay times (τ < 3s) was used to obtain a linear fit. Diffusivity were confirmed by estimating at τ = 1s.

We then repeated the procedure and extracted tracer diffusivities in the 10 wt% mucin solution using the ensemble averaged MSD’s. As seen from the figure, tracer diffusivity in the bare mucin solution also follows the expected Stokes-Einstein relationship. Since the tracer size and temperature are known, this data can be used to extract the effective viscosity of the solution. We find consistent values when data from tracers with diameter 2.11 *µ*m and smaller are used. Larger tracers however are found to have significantly lower than predicted diffusivity values (assuming the ambient medium viscosity is the same for all tracers). In subsequent sections, we will investigate this result by examining the probability distribution of individual mean square tracer displacements to identify why the mobility may be reduced.

### C. Diffusivity in rock-dust laden mucin formulations is larger than in the bare mucin

We next investigated if the addition of anti-caking agents and dust affects tracer diffusion. Again, the data for ⟨ MSD ⟩ was consistent with a *γ* ≈ 1 exponent as the best fit suggesting very weak viscoelastic effects and suggesting that tracers felt a fluidic albeit complex rheological environment. Figure 6 (b) is a plot of the diffusivities for various formulations and for increasing tracer sizes. We find that for all agents tested, the diffusivity of tracer particles (at fixed *a*) increases compared to the 10 wt% control mucin solution.

The diffusivity measured in this text differs from that estimated in [15], where inclusion of anti-caking agents reduce overall diffusivity. Consistently however, in their study and in ours, we find that diffusivity (of tracers in our study) in mucin with MTRD solutions is higher than in mucin with UCRD or SiO_2_. This is not surprising for a variety of reasons. First, the diffusivity in [15] is obtained from swelling experiments; that is the diffusive flow of water (the tracers here being of molecular dimensions) through a gel-like network is studied. Our samples do not form gels but are solutions. Second, our methodology hinges on tracer motion, and explores local properties as opposed to the global properties.

To identify possible mechanisms that impact thermally driven transport in the dust-laden mucin samples, we consider biochemical effects and possible effects of rock dust on mucin network structure and morphology. Anticaking agents used in the mining industry are usually hydrophobic [44] to prevent the swelling of dust that leads to caking after moisture evaporation. Electrostatic interactions between the hydrophobic dust molecules and water in mucin solutions may allow efficient transport of the tracers by preventing mucin chains from entangling together. Calcium present in the solutions due to release from the dust particles may act as crosslinkers slowing mucin swelling rates, and therefore reducing diffusivity when measured through swelling kinetics [45, 46]. Concomitantly, diffusivity of particles may be increased due to electrostatic effects caused by calcium leaking from dust coatings [47]. Rock dust in the medium secretes calcium ions which are charged and can be highly mobile throughout the mucin network. These charges may then be picked up by tracers. The Debye screening of the charged tracer may prevent them from physically interacting with mucin chains, enabling faster movement through the medium. Lielieg *et al*. [47] find that while there is negligible change in the diffusivity of 1 *µ*m neutral PEG tracers in mucin, there is a small increase at high salt concentrations that may account for our increases in diffusivity in mucin with rock dust solutions.

From a consideration of these prior studies, we suggest that two competing effects may be playing a role here: 1) crosslinking of mucin chains (due to cationic calcium ions leaching into the mucin from added particles or anti-caking agents) which lead to reduced mobility of microscopic tracers, and 2) electrostatic interactions between charged tracers (that pick up charges as they move in the mucin) and entangled and dangling chains in the loosely connected mucin network that increase the motion of tracers relative to the environment.

### D. Tracking tracer squared displacements highlights effects of heterogeneity and transient trapping

In isotropic, isothermal, viscous media, the diffusion of a spherical tracer with a diffusivity *D* can be quantified either by the probability distribution of the displacement magnitude, or equivalently by the distribution of the squared displacement. Following a number *N*_τ_ of these tracers for a time interval τ (the time from start to the end of the observation period with the displacement calculated over this period), the probability density of displacement Δ^2^ follows [40],

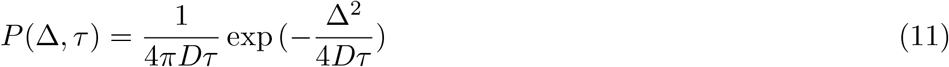

valid so long as the measurement time interval is much smaller than *L*^2^*/*4*D* where *L* is the overall dimension of the imaged space. The probability of the signed displacement has mean zero and a variance that grows with time. In Figure 3(a), we validated this result for tracers in DI water (we plot the displacement histogram distribution in the figure, not the squared displacement). When tracers move in non-ideal environments where their effective transport properties are spatiotemporally dependent - such as due to interactions with polymer networks such as trapping or caging, local heterogeneities, or tracer-tracer interactions, the distribution of displacements is expected to deviate from the ideal form. Examination of the measured distributions will there provide information about the properties of the environment around the moving tracers.

We estimated particle histograms (discrete distributions) for various lag times and for different tracer sizes; this data was then used to discern the effect of the medium on the dispersion of the particles. In generating such histograms from particle trajectories in the different mucin formulations, we made sure that the tracers tracked were in the same local environment (roughly the dimension of the imaging area) so that the distribution obtained is a measure of non ideal effects such as transient trapping or viscoelasticity, manifest in the imaged region. Even then, in several instances, we identified tracer trajectories averaged MSD that were very different suggesting impact due to micro-scale heterogeneities.

Before investigating histograms for tracers moving in the various mucin solutions, we studied how particles moved in viscoelastic CMC. This provides a baseline for understanding effects of polymer networks and viscoelasticity on tracers but without the complicating effects of rock dust. Figure 7 shows the histograms obtained for a 1 *µ*m tracer at concentrations from 0.5% (weakly elastic fluid) to 2% (strongly viscoelastic). Three different lag times are shown and all cases have the same number of trajectories. We first note that the mean displacement of the tracers tracked for fixed τ decreases with concentration as expected. The histograms for lower concentrations are as expected with a clear peak that decreases with lag time, accompanied by a spread in values (dispersion). The 2% concentration data shows significant differences however. First we find that the peak for τ = 1s to be higher than for the small and large lag times. Second, comparing the distributions for τ = 0.1s, we find that asymmetric distribution with a long tail for the 2% concentration. This is not because some tracers have undergone mean square displacements higher than expected; rather, this is because a significant number of tracers are being hindered in their movement. This hindered motion seems to relax as the lag time increases to τ = 10s.

**FIG. 7.**
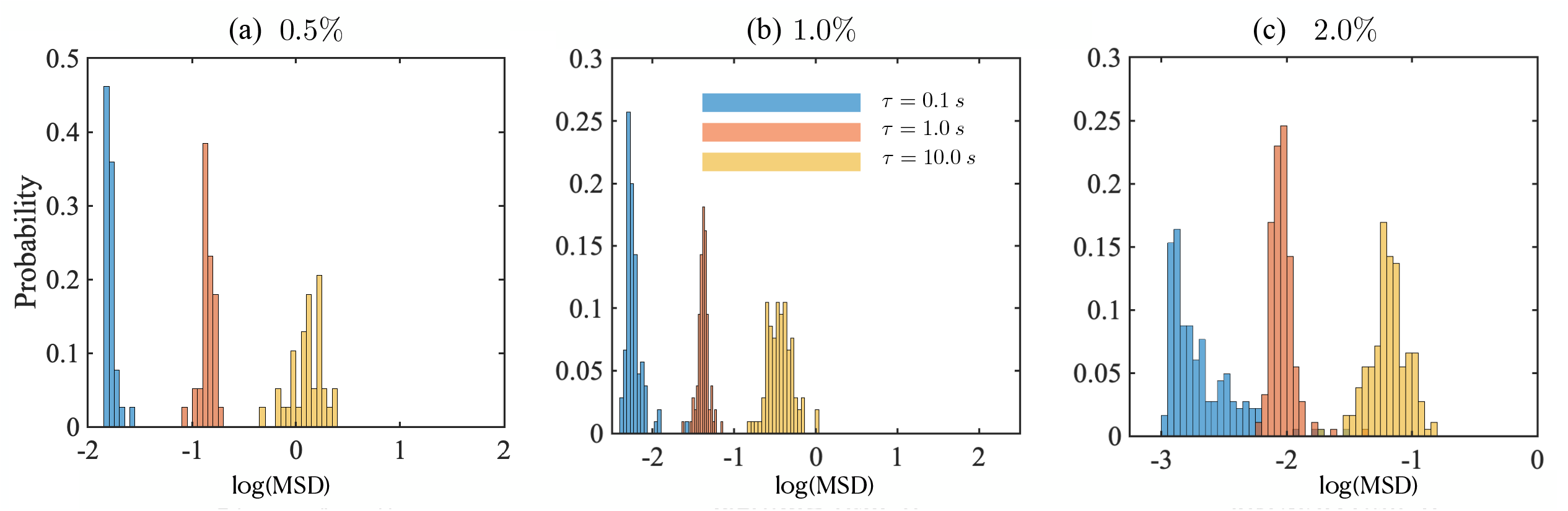
Histograms denoting the discrete probability distribution of (trajectory averaged) mean square displacements (MSD) for 1 *µ*m tracer particles in 250 kD CMC solution at three delay times τ = 0.1, 1, 10 seconds. We show results for three CMC concentrations from 0.5% to 2%.

A similar analysis conducted for the control 10 wt% mucin solution without any added rock-dust is shown in Figure 8. Within each tile, the number of trajectories (particles) followed is constant. We focus on some interesting trends. First, starting from the smallest tracer (*a* = 0.5 *µ*m), we observe that the MSD values typically decrease with increasing tracer diameter for all values of τ as expected. However that except for the 1 *µ*m tracer particle, there is no clear reduction in the peak value with increasing τ. This is especially surprising for the smallest tracer size.

**FIG. 8.**
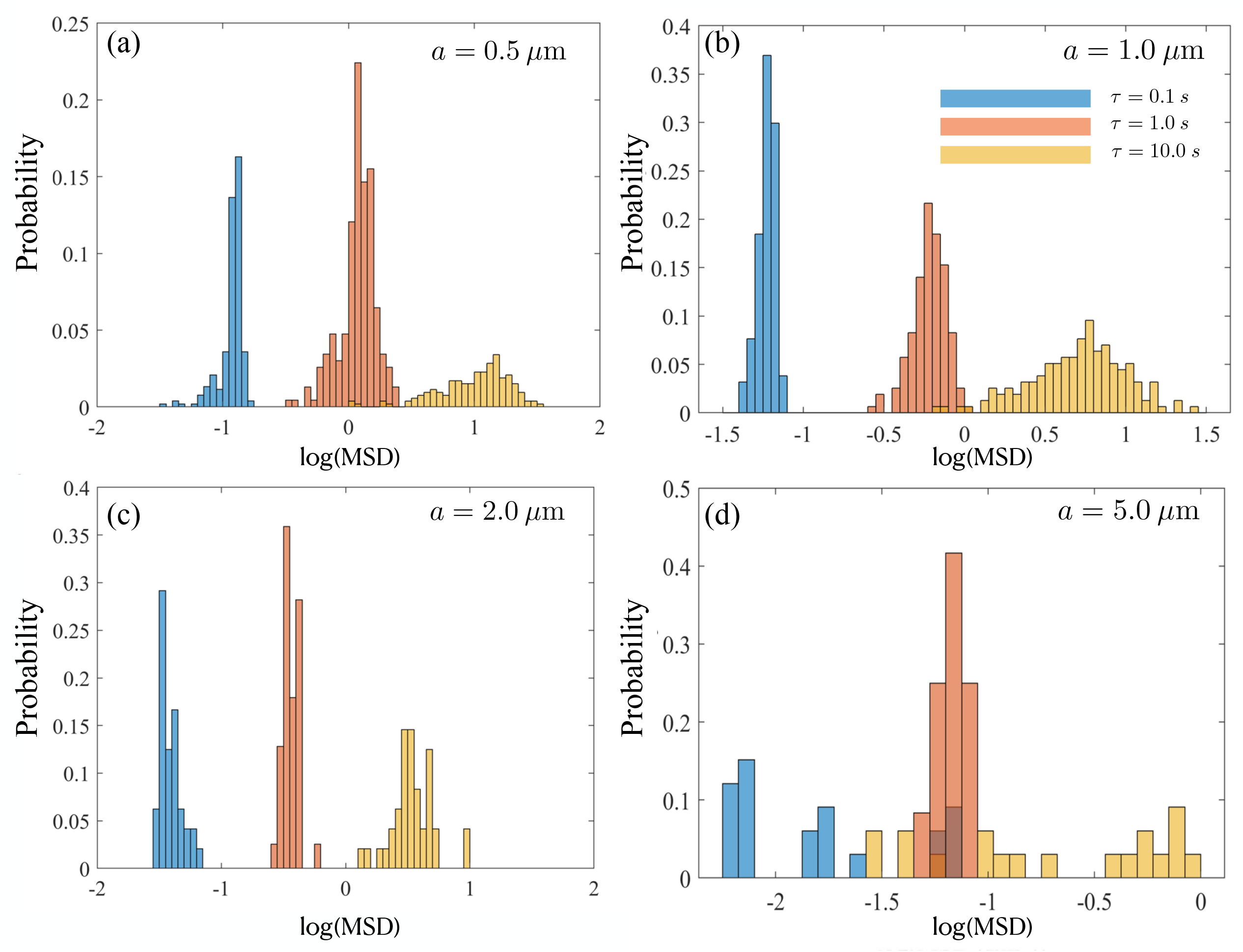
Histograms denoting the probability distribution of trajectory-averaged MSD, shown for the 10 wt% control mucin solution at delay times τ = 0.1, 1, 10 seconds. We show data for *a* = (a) 0.5 *µ*m, (b) 1 *µ*m, (c) 2 *µ*m, and (d) 5 *µ*m. For the largest tracer size of 5.0 *µ*m, the variation in values is larger compared to smaller tracer sizes. The number of trajectories analyzed are (a) *N* = 270, (b) *N* = 157, (c) *N* = 76, and (d) *N* = 20. Images were taken on Zeiss 200M Axiovert microscope with 40x/NA 0.75 objective at 30 fps, 30 ms exposure time.

Secondly, compared to small tracers with *a* = 0.5 and *a* = 1.0 *µ*m, the histogram distribution for large tracers (at each value of τ) is significantly distorted and asymmetric. For instance the 2.0 *µ*m tracer shows evidence of hindered mobility for small lag times. At τ = 10 s, a significant fraction of the tracers display a reduced value of the MSD suggesting short-time trapping and release as moving tracer sample and interact with the polymers in solution. While we have not characterized the typical mesh sizes of the loosely entangled polymers in the mucin solution, relaxation of moderately sized particles suggests that caging effects due to mucin-polymer interactions are transient. For the largest tracer examined (*a* = 5 *µ*m), the histograms are consistent with the low values of the diffusivity we found. We also see significant spread at long delay times (τ = 10 s). As mentioned previously, we found that large tracers within the same mucin sub-region tended to have a wide diversity in MSD values suggesting that micro-scale heterogeneities of the embedding medium are also contributing to the wide range in MSD values.

We next studied histograms of MSD values for tracers moving in the dust-laden mucin solutions. Each type of rock dust has a distribution of sizes (see Table 1) with mean that is in the micron range with a substantially large fraction larger than the size of the tracers we have used. Intuitively, we expect that the addition of rock dust increases the effective mean viscosity of the mucin solution. As discussed earlier in Section §3.3, this does not readily translate to reduced diffusivities compared to the 10 wt% control mucin. Tracers moving in UCRD laden solutions have slightly higher diffusivity values compared to MTRD and SiO_2_ dust, except for the smallest tracer where they are roughly the same. The rock dust size distribution (Table 1) indicates that the smallest mean size ∼3.4 *µ*m corresponds to SiO_2_ dust for which we expect inter-dust mucin filled voids to be smaller. Tracers then have to move through reduced spaces, and furthermore interact more frequently with the rock dust. These effects may explain why the average diffusivity value obtained from ⟨ MSD ⟩ (averaging over multiple particles) is lower for the 10% mucin with SiO_2_. Indeed, we find from Figure 9 that the MSD histograms for mucin-MTRD and mucin-UCRD are shifted more to the right than for mucin-S. Further quantification is unfortunately not possible since we do not know the exact distribution of dust sizes. Furthermore, mucin solutions with rock dust showed evidence of aggregation and clumping. We also note from Figure 9 (a) that the histogram for mucin-MTRD (Mucin-CD in the figure) is symmetric compared to those for mucin-UCRD and mucin-S. To understand how this feature changes with *a* and τ, we analyze the appropriate MSD histograms (Figure 10). As τ increases with fixed *a*, the peak in frequency goes down and the characteristic MSD value increases. As *a* increases, the log(MSD) histogram plots shift to lower magnitudes as expected but show evidence of the mobility of tracers being hindered for small τ. For instance in (b), we see pronounced fore-aft asymmetry for τ = 0.1 s but not as much for τ = 10 s. The largest tracer results show significant scatter in values, and distributions skewed by mobility impaired tracers.

**TABLE 1.**

| **TABLE 1** Rock dust type | Particle size details |
| --- | --- |
| MineBrite™-G (UCRD) | 97% $< 74 \mu\text{m}$, 24% $< 10 \mu\text{m}$ |
| ImerCoal™ - Moisture Tolerance Rock Dust (MTRD) | Mean: $19.5 \mu\text{m}$ |
| MIN-U-SIL® 10 Ground Silica ($\text{SiO}_2$) | Mean: $3.4 \mu\text{m}$ |

**FIG. 9.**
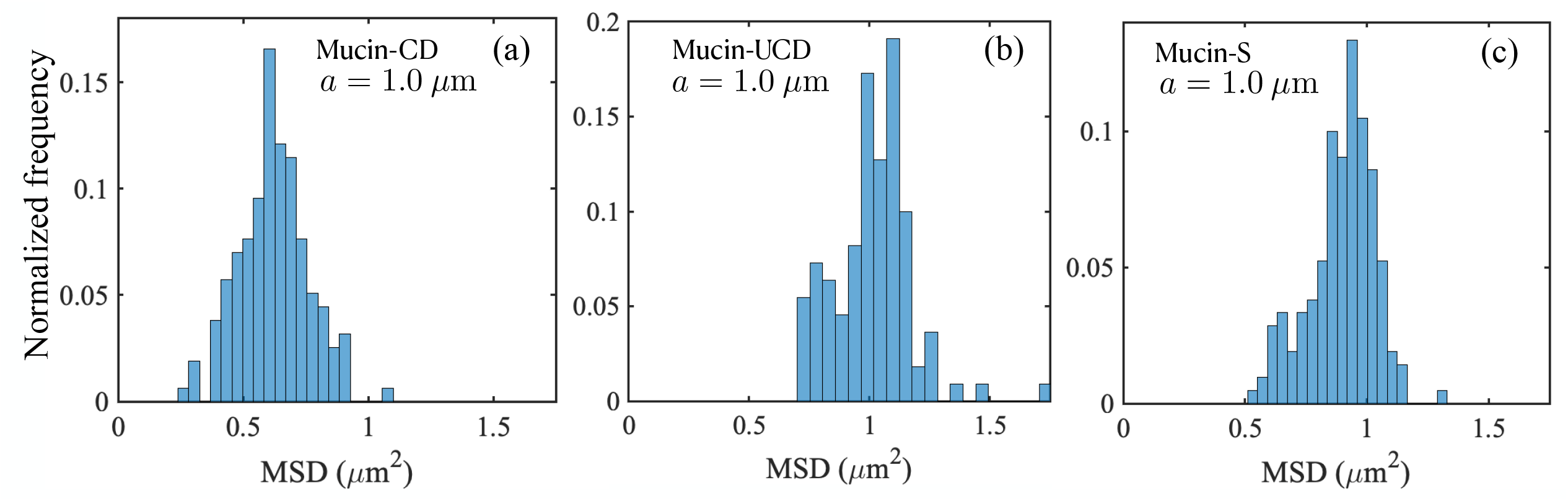
Histograms obtained by analyzing the squared displacements of 1 *µ*m tracers in dust-laden mucin samples estimated after a lag time τ =0.1 s. **(a)** Histograms for mucin with MCRD, with number of samples *N* = 109. **(b)** Histograms for mucin with UCRD, with number of samples *N* = 173. **(c)** Histograms for mucin with SiO2, with *N* = 157. MCRD with mucin shows lower values of measured MSDs compared to UCRD and SiO2.

**FIG. 10.**
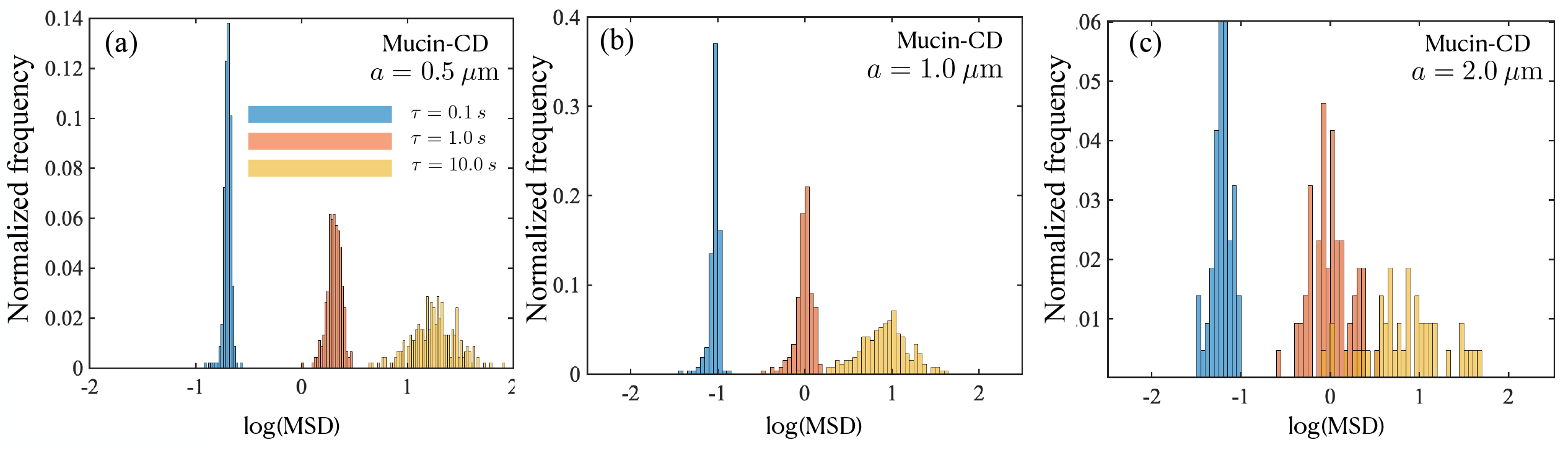
Histograms for mucin samples with MCRD. The tracer vary from 0.5-2.11 *µ*m in diameter, and three delay times are shown (τ = 0.1, 1.0 and 10.0 s (blue, orange, yellow, respectively)).

## IV. CONCLUSIONS

The thermally driven diffusion and dispersion of particulates in mucin is relevant to understanding how inhaled aerosols and dust may penetrate the mucus layer and be transported through it. Motivated by the typical sizes of rock dust encountered in mining environments, and similar environmental aerosolized pollutants, we focus on sub-micron to micron tracers larger than around 500 nm. Here, we demonstrate that high-speed particle tracking combined with passive microrheology provides a rigorous tool to study the motion of these particulates, as well as quantify the complex rheology and viscoelasticity the embedding medium. We validate our methods and confirm their accuracy by estimating the diffusivity of spherical tracers in DI water, displacement histograms, and velocity auto-correlation function and validating these with expected theoretical results. We then utilized our methods to determine the motion of 0.5-5.0 *µ*m spherical tracers in 10% mucin solutions with and without suspended rock-dust. The particle size dependence of the calculated diffusivity for all types of mucin solutions follows the Stokes-Einstein relationship for particles ∼1 *µ*m and smaller. Larger ∼3-5*µ*m particles show evidence of sub-diffusive behavior. This is confirmed by analysis of the probability distribution of the mean square displacements. Taken together, our analysis of the MSD distributions suggest that heterogeneity, trapping, and electrostatic effects impact tracer transport in mucin, especially for large tracer particles. Our results motivate further studies on how heterogeneity, transient trapping due to tracer-network interactions, electrostatic effects, and viscoelasticity may impact particle transport in mucin solutions and gels.

